# H3K27me3 chromatin heterogeneity reveals variable cell responses to estrogen and endocrine treatment

**DOI:** 10.1101/2025.09.18.677184

**Authors:** Fleur Chapus, Christopher R Day, Laura G Kammel, Pelin Yasar, Brian Bennett, Erica Scappini, Charles Tucker, Maria Sifre, Celyn Bregio, Joseph Rodriguez

## Abstract

Gene expression heterogeneity generates subpopulations of tumor cells that can evade therapeutic pressure. This heterogeneity has been observed in both primary Estrogen Receptor alpha positive (ERα+) breast tumors and cell lines. Therefore, understanding the mechanisms regulating expression heterogeneity is critical towards developing effective therapies. A key contributor to gene expression variability is the stochastic nature of transcription. Transcription occurs in a probabilistic, burst-like manner, in which gene activation occurs intermittently, producing RNA in pulses and interspersed with transcriptional off-periods. The estrogen-responsive gene *TFF1* is expressed in the majority of ERα⁺ breast tumors and exemplifies such heterogeneity, with transcriptional inactivity ranging from minutes to several days. Here, we identify the molecular mechanism underlying the wide range in TFF1 expression by analyzing cells sorted based on their TFF1 activity levels. We observed that TFF1 inactive (TFF1^low^) cells exhibit a repressive chromatin state marked by H3K27me3 at the TFF1 promoter and enhancer. Despite global similarity in ERα binding, occupancy at the *TFF1* regulatory elements was selectively reduced in TFF1^low^ cells, resulting in fewer active alleles and diminished transcriptional bursting frequency. Conversely, TFF1^high^ cells exhibited more active TFF1 alleles and hyperbursting. These cells also retained sensitivity to endocrine therapy, while TFF1^low^ cells displayed reduced drug responsiveness. Genome-wide, differentially enriched H3K27me3 regions correlated with variable expression of estrogen-responsive genes, highlighting a broader regulatory mechanism that links chromatin state to expression variability. Together, our findings establish how repressive chromatin dynamics contribute to gene expression heterogeneity and endocrine resistance in ERα⁺ breast cancer.

**Graphical abstract:** 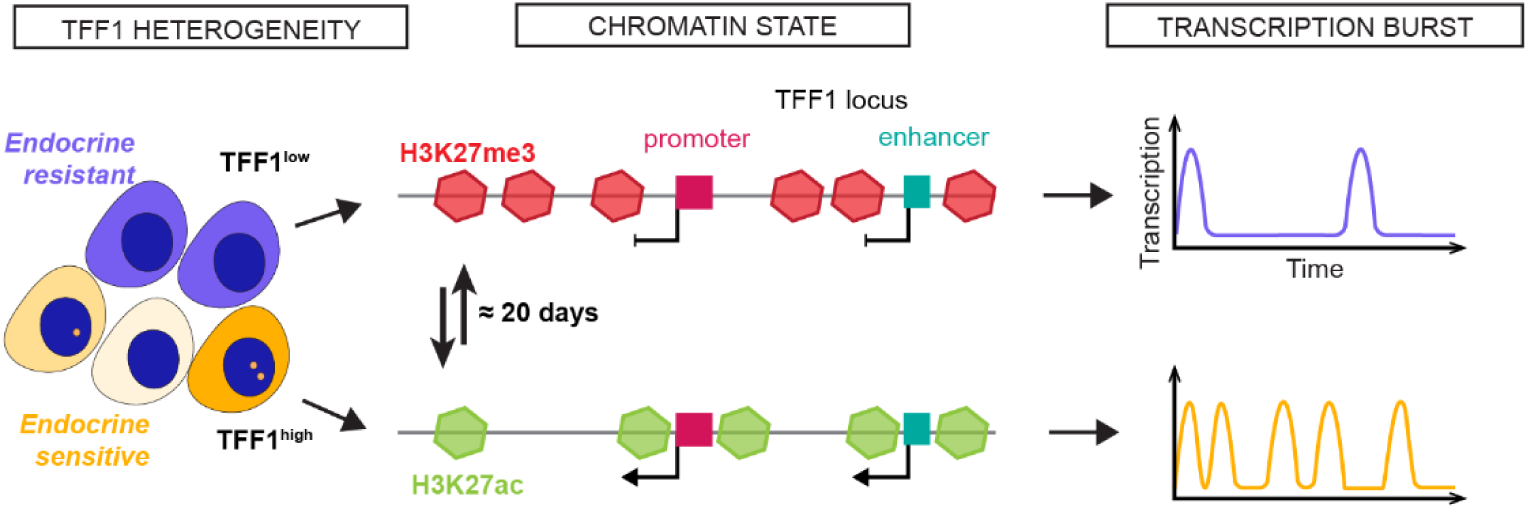

## Introduction

Intra-tumoral heterogeneity arises from differences in gene expression among tumor cells, driven by both genetic and epigenetic processes^1^. This variability in gene expression results in differential protein concentrations among individual cancer cells, leading to functional diversity within the tumor cell population. Hence, differences in gene expression are directly associated with cancer progression, as subpopulations of cells may acquire resistance or metastatic traits through non-genetic mechanisms^2–4^, posing challenges for treatment strategies.

One source of non-genetic variability is stochastic transcriptional activity, where individual cells experience alternating periods of active gene expression or inactivity at individual gene alleles^5,6^. This phenomenon is known as transcriptional bursting and contributes to gene expression variability by randomly switching genes between high and low transcriptional states. In Estrogen receptor alpha (ERα)-positive breast, the response to estrogen generates variability in gene expression. ERα+ breast cancer accounts for approximately 80% of invasive breast cancers and is generally associated with a favorable prognosis when treated with endocrine therapies^7^. However, resistance to these therapies, such as selective estrogen receptor modulators (SERMs) and down-regulators (SERDs), can emerge over time, highlighting the need to address intra-tumoral heterogeneity to improve therapeutic strategies ^8–10^.

Trefoil factor 1 (TFF1) is an estrogen-regulated gene and ectopically expressed in 70% of ERα+ breast tumors^11^. As previously demonstrated, highly variable levels of TFF1 RNA are expressed within tumor cell populations^12^. The luminal breast cancer cell line MCF7 was shown to be a good model for this variability as TFF1 transcription in response to estrogen (17β-estradiol, E2) stimulation is marked by a pronounced heterogeneity^12^. The variability of TFF1 expression is largely attributable to an uncharacterized deep repressive state. This state is marked by prolonged transcriptional inactivity that can persist from minutes to several days, even under activating conditions^12^. Interestingly, high TFF1 expression correlates with better prognosis^13,14^ and improved overall survival^15,16^. However, the mechanisms dictating TFF1 transcriptional dynamics and the implications of its expression variability in breast cancer progression remain to be elucidated.

Possible mechanisms underlying this deep repression at estrogen-responsive genes involve chromatin-modifying complexes such as Polycomb group proteins and histone deacetylases (HDACs). Both Polycomb repressive complexes (PRC1 and PRC2) and HDACs are known to maintain long-term gene silencing through histone modifications, primarily H3K27 trimethylation and deacetylation^17^, respectively. These repressive marks contribute to chromatin compaction and reduced accessibility of transcription factors at target gene loci. Previous studies have implicated both Polycomb and HDACs in regulating estrogen-responsive genes^18–21^, suggesting that they may stabilize the deep repressed state observed at TFF1 and other ERα targets. Therefore, these epigenetic regulators represent candidate mechanisms in maintaining transcriptional silencing and driving heterogeneity in ERα+ breast cancer cells.

Here, we investigate the molecular mechanisms of TFF1 transcriptional heterogeneity, and how its transcriptional dynamics contribute to endocrine resistance. For this study, we used a TFF1 protein-based cell sorting strategy that separates cells by TFF1 expression. The chromatin landscape and transcriptional activity of high and low TFF1-expressing cells were profiled using genomics approaches, single molecule RNA FISH (smFISH), and live cell imaging. We discover a heterogeneous chromatin landscape at the TFF1 promoter and enhancer which is marked by repressive chromatin in TFF1^low^ expressing cells. In contrast, TFF1^high^ cells exhibit increased ERα binding and TFF1 transcriptional burst initiation. We generalize these findings to other estrogen-responsive genes and link these heterogeneous chromatin states to differential responses to cancer therapeutics.

## Results

### TFF1 heterogeneity is dictated by a heterogeneous chromatin landscape at the promoter and enhancer

70% of ERα+ breast cancers express the estrogen-responsive TFF1 gene, with the level of expression varying widely among tumors and within cells in those tumors^14^. To identify the mechanism underlying this heterogeneity, we hypothesized that sorting cells by their protein expression would permit genomic characterization of the chromatin landscape for different activity levels. To determine if we could use TFF1 protein to enrich for cells with high and low RNA levels, we first used protein immunocytochemistry combined with single-molecule RNA FISH (ImmunoFISH) in MCF7 cells treated with estrogen (17β-estradiol, E2) for 48 hours. We observed heterogeneous expression for both the protein and mRNA signals (Figure 1a). We next divided the cells into two groups based on TFF1 immunofluorescence intensity: the top 20% (« Top20% ») and the remaining 80% (« Rest »). For each group, mRNA counts per cell were quantified from smFISH signals. We observed a significant enrichment in TFF1 mRNA counts per cell in the high-expressing cells compared to the rest of the population (Figure 1b). These data indicated that TFF1 protein could be used to select for cells of different activities. We next established a fluorescence-activated cell sorting (FACS) strategy using immunofluorescence labeling of the TFF1 protein in live MCF7 cells. Cells were cultured in hormone-depleted medium for 72 hours before adding 1nM E2 for 48 hours. We then isolated the top 20% of TFF1 expressing cells (TFF1^high^) and the lowest TFF1-expressing cells (TFF1^low^), keeping an approximately 15% margin between the 2 cell populations to capture the two ends of the TFF1 expression distribution (Figure 1c; Supp. Figure 1a). We first asked whether transcriptional activity could explain the differential RNA output observed between TFF1^high^ and TFF1^low^ cells. We performed Nascent RNA sequencing (Nascent-seq) by isolating newly synthesized chromatin-bound RNA using a high salt lysis buffer^22^(Supp. Figure 1b). Differential gene expression analysis using DESeq2 confirmed that TFF1 was the most differentially expressed gene (DEG) in TFF1^high^ compared to TFF1^low^ cells (Supp. Figure 1c), validating TFF1 protein-based sorting as a relevant method for further investigating TFF1 heterogeneity. Cells sorted into TFF1^high^ and TFF1^low^ populations displayed correspondingly high and low levels of TFF1 promoter activity (Log_2_FC = 1.413; adj p = 6.92E-05) (Figure 1d). Strikingly, enhancer RNA transcription was also higher in TFF1^high^ cells than in TFF1^low^ cells, suggesting co-regulation of the TFF1 promoter and enhancer RNA transcription. Moreover, the reduced transcription spans the entire TFF1 locus in TFF1^low^ cells and indicates the existence of a synchronous, deeply repressive state at both TFF1 promoter and enhancer regions of all alleles.

**Figure 1.**
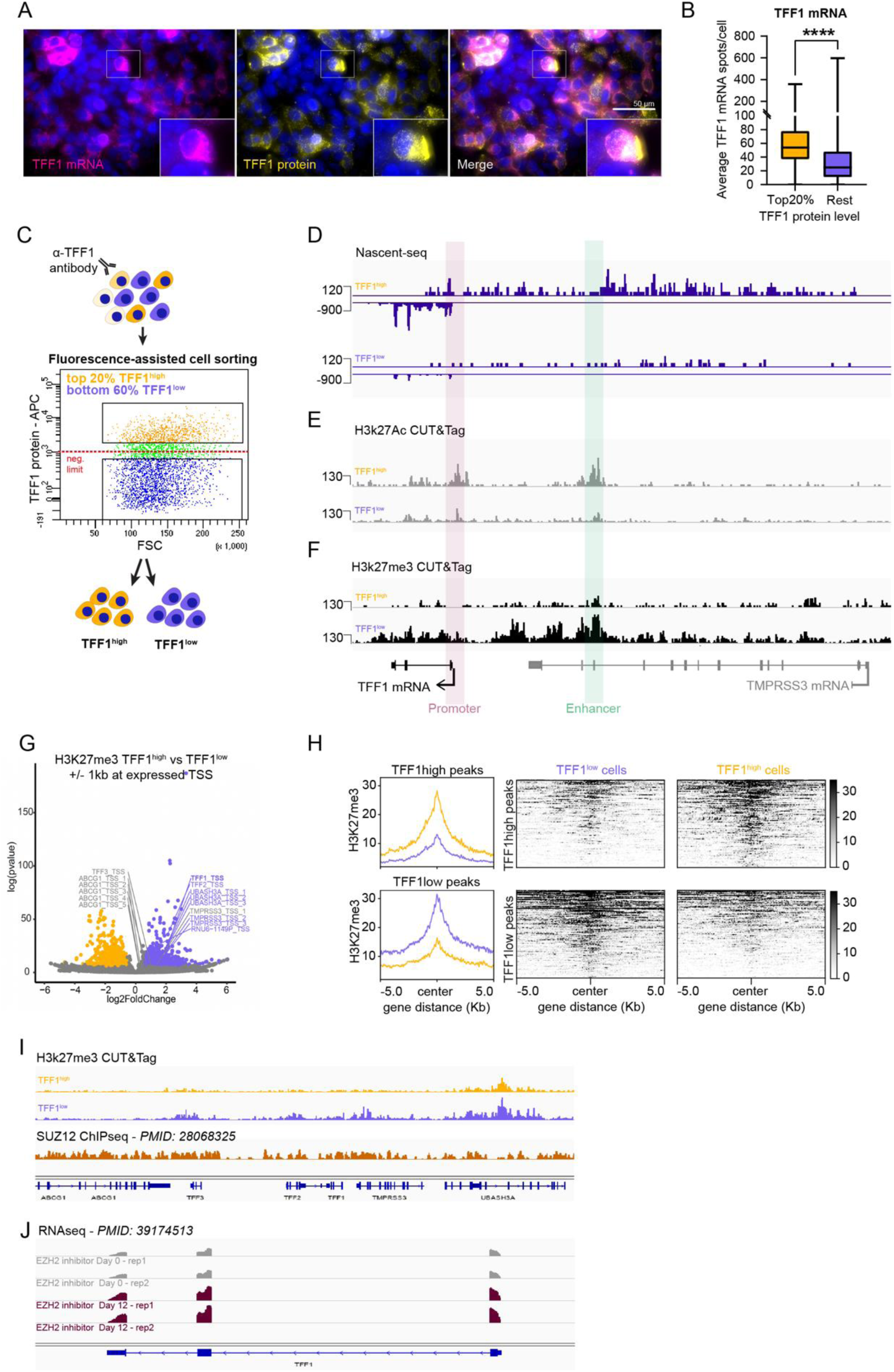
TFF1 heterogeneity is dictated by a heterogenous chromatin landscape at the promoter and enhancer MCF7 cells have been treated with 1 nM of E2 during 48 hours. A. Immuno-smFISH image of heterogenous TFF1 mRNA (magenta) and protein (yellow) expression in MCF7 cells. Scale bar = 50 μm B. Immuno-smFISH quantification: Top 20% of TFF1 protein expressing cells contain significantly more mRNA than the rest of population. C. Schematic illustrating cell sorting strategy. D. Genome browser view of Nascent-seq of TFF1^high^ and TFF1^low^ cells at TFF1 gene and enhancer locus. E. Genome browser view of H3K27ac CUT&Tag at TFF1 gene and enhancer locus in TFF1^high^ and TFF1^low^ cells. F. Genome browser view of H3K27me3 CUT&Tag at TFF1 gene and enhancer locus in TFF1^high^ and TFF1^low^ cells. G. Volcano plot of H3K27me3 differential promoters in TFF1^high^ and TFF1^low^ cells show multiple differentially regions in TFF1 local neighborhood. H. Metaplots and heatmaps of H3K27me3 CUT&Tag peaks enriched in TFF1^high^ and TFF1^low^ cells. I. Genome browser view of H3K27me3 CUT&Tag and public MCF7 SUZ12 ChIP-seq data in TFF1 local neighborhood. J. Genome browser illustration of public MCF7 data showing EZH2 inhibition increases TFF1 expression after 12 days.

Given that transcriptional activity is influenced by the epigenetic status at the promoter and enhancer regulatory regions^23^ we next determined whether the chromatin landscape also reflected the different transcriptional states observed at the TFF1 locus. We examined the deposition of the active chromatin mark H3K27ac and the repressive H3K27me3 histone mark using CUT&Tag in TFF1^high^ and TFF1^low^ sorted cell populations. As expected, in TFF1^high^ cells, active transcription corresponded to increased H3K27ac deposition at the TFF1 locus (Figure 1e). Strikingly, the absence of transcription at the TFF1 locus in TFF1^low^ cells co-occurred with significantly increased H3K27me3 deposition (Figure 1f) at both the promoter and enhancer. We next quantified global changes in H3K27me3 by using a +/-1-kb window around 87,837 expressed Transcription Start Sites (TSS) identified from publicly available Start-seq data in MCF7 cells^24^. We observed that the vast majority of TSSs had no differential H3k27me3 deposition between TFF1^high^ and TFF1^low^ cell populations. However, we did observe 1063 TSSs (1.2% of all expressed TSSs) with increased H3K27me3 and 1105 TSS (1.25% of all expressed TSSs) with decreased H3K27me3 (adj p <0.05) (Figure 1g,h). Importantly, in addition to TFF1, the TSSs for TFF2, TMPRSS3 and UBASH3A were among the 1063 TSSs with significantly greater H3K27me3 deposition in TFF1^low^ cells (Figure 1g). These genes neighbor TFF1 and form a sub-topologically associated domain (subTAD) spanning approximately 150kb^25^ (Figure 1i). Given that PRC2 catalyzes the trimethylation of H3K27me3, we asked whether this locus was also bound by the PRC2 subunit SUZ12 in MCF7 cells using publicly available data^26^. Indeed, we observed SUZ12 deposition across the locus (Figure 1i). To interrogate the potential functional role of PRC2 in repressing TFF1 transcriptional activity, we took advantage of a publicly available RNAseq dataset^27^ when inhibiting the catalytic subunit of PRC2, EZH2 in MCF7 cells. Interestingly, the inhibition of EZH2 during 12 days increased the level of TFF1 mRNA compared to the non-treated condition (Figure 1j). Taken together, these data suggest that PRC2 plays a role in cells with silenced TFF1 alleles, and that a subset of TSSs exhibit differential H3K27me3 enrichment between TFF1^high^ and TFF1^low^ cells^26^. We also identify H3K27me3 repressive chromatin as the molecular basis of the deep repressive state for TFF1.

### Repressive chromatin landscape prevents ERα binding at the TFF1 locus

Binding of the transcription factor ERα at the promoter and enhancer is critical for TFF1 activation^28^. To determine if the differential H3K27me3 chromatin affects the ability of ERα to bind to TFF1 regulatory elements, we profiled ERα binding in TFF1^high^ and TFF1^low^ sorted populations using CUT&Tag. In total, we obtained 22,929 ERα peaks corresponding to 10,982 peaks in the TFF1^high^ cells and 11,947 peaks in the TFF1^low^ cells. We observed a high degree of ERα binding similarity between the two sorted populations. Only 1.9% and 2% of the peaks were differential peaks for the TFF1^high^ and TFF1^low^ datasets, respectively (Figure 2a,b; Supp. Figure 2a-e). These data indicate that the majority of called ERα peaks are similar in binding magnitude. However, ERα binding was significantly higher at the EREs of the TFF1 promoter and enhancer in TFF1^high^ population than in the TFF1^low^ population (enhancer adj p = 4.15e-7; promoter adj p = 2.72e-6) (Figure 2c). This data suggests that the repressive chromatin environment at TFF1 locus inhibits ERα binding. Taken together, we have classified TFF1 heterogeneity into different activity states and observed that the deeply repressive state at the TFF1 locus was due to repressive H3K27me3 chromatin landscape and decreased ERα binding.

**Figure 2.**
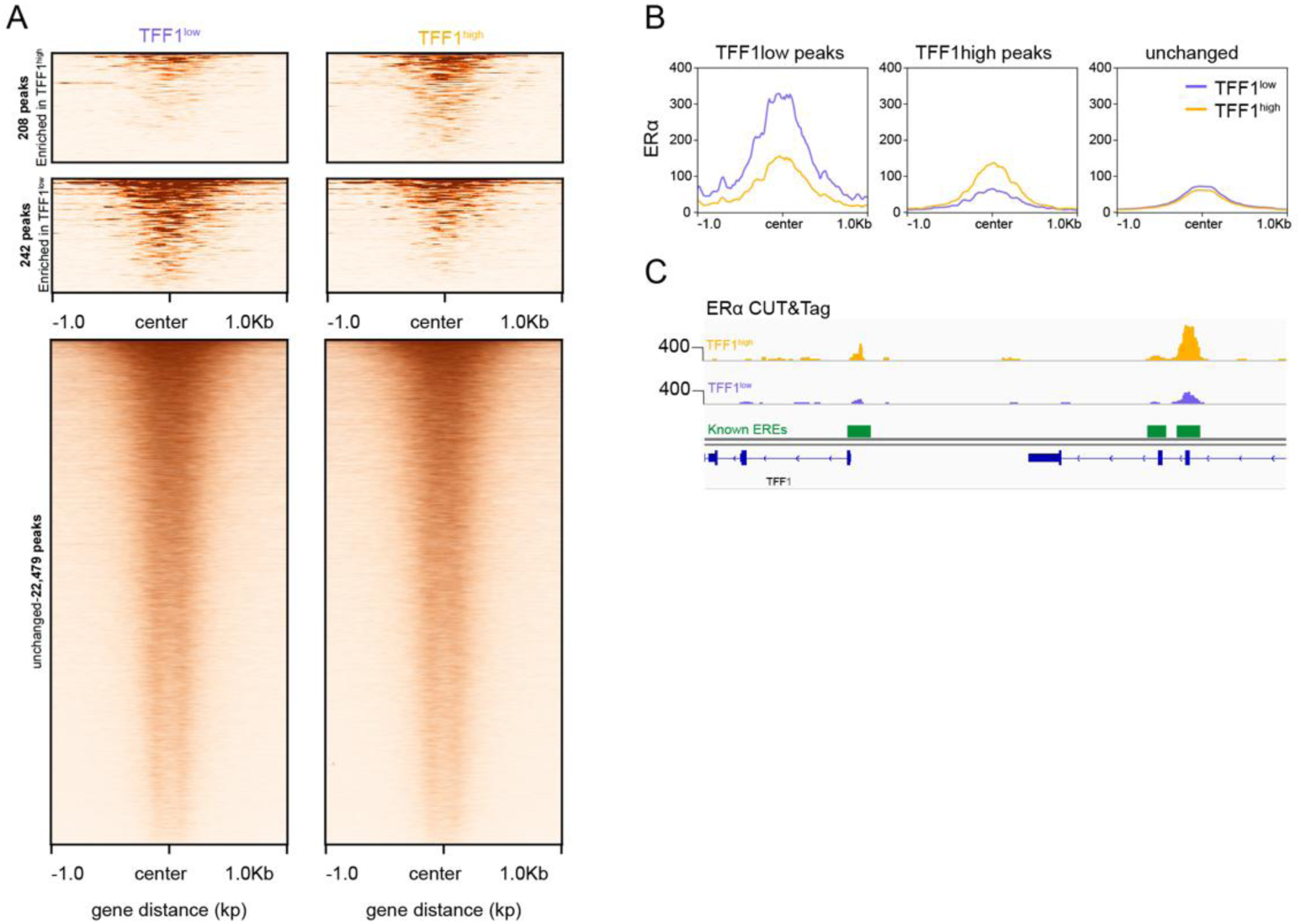
Repressive chromatin limits ERα binding at TFF1 locus. A. Filtered heatmaps of ERα binding in TFF1^high^ and TFF1^low^ cells shows that the majority of ERα sites are unchanged. B. Metaplots of ERα binding enriched in TFF1^high^ cells and TFF1^low^ cells and unchanged peaks. C. Genome browser view of ERα CUT&Tag shows that ERα binding is significantly higher at the TFF1 promoter an enhancer in TFF1^high^ cells compared to TFF1^low^ cells.

### High TFF1 expressing cells exhibit hyper bursting

To gain a deeper mechanistic understanding of how the differential chromatin landscape and ERα binding at TFF1 regulatory regions affect transcriptional activity, we characterized the bursting dynamics of the TFF1 gene in TFF1^high^ and TFF1^low^ populations. We took advantage of a TFF1-MS2 MCF7 cell line expressing 24 repeats of the MS2 RNA stem loops endogenously integrated into the 3’UTR of TFF1 gene^12^. This cell line enables the real-time visualization of TFF1 transcription via the emission of a fluorescent signal at the site of transcription when the promoter is transcriptionally active^12^ (Figure 3a). TFF1-MS2 cells were sorted for different TFF1 protein levels as described above and imaged the day after sorting for 14.2 hours to capture bursting kinetics^12^ (Figure 3b; Supp. Figure 2a,b; movies 1 and 2) We quantified the transcriptional behavior at the TFF1 promoter by tracking the fluorescence intensity at the site of transcription over time and applied a Hidden Markov Model (HMM) to define active and inactive states (Figure 3c,d). For each TFF1^high^ and TFF1^low^ population, a total of 50 alleles were tracked, and the median active time (Figure 3e,f) and the median inactive time (Figure 3g,h) were calculated. Strikingly, while TFF1 median active time remained identical between the two populations (TFF1^high^ = 6.67 mins; TFF1^low^ = 6.67 mins), the periods of inactivity significantly increased in TFF1^low^ cells (TFF1^high^ = 43.33 mins; TFF1^low^ = 63.33 mins). These data suggest that differential chromatin and ERα binding regulate bursting kinetics at the TFF1 promoter by promoting more frequent transcription initiation in TFF1^high^ cells compared to their TFF1^low^ counterparts.

**Figure 3.**
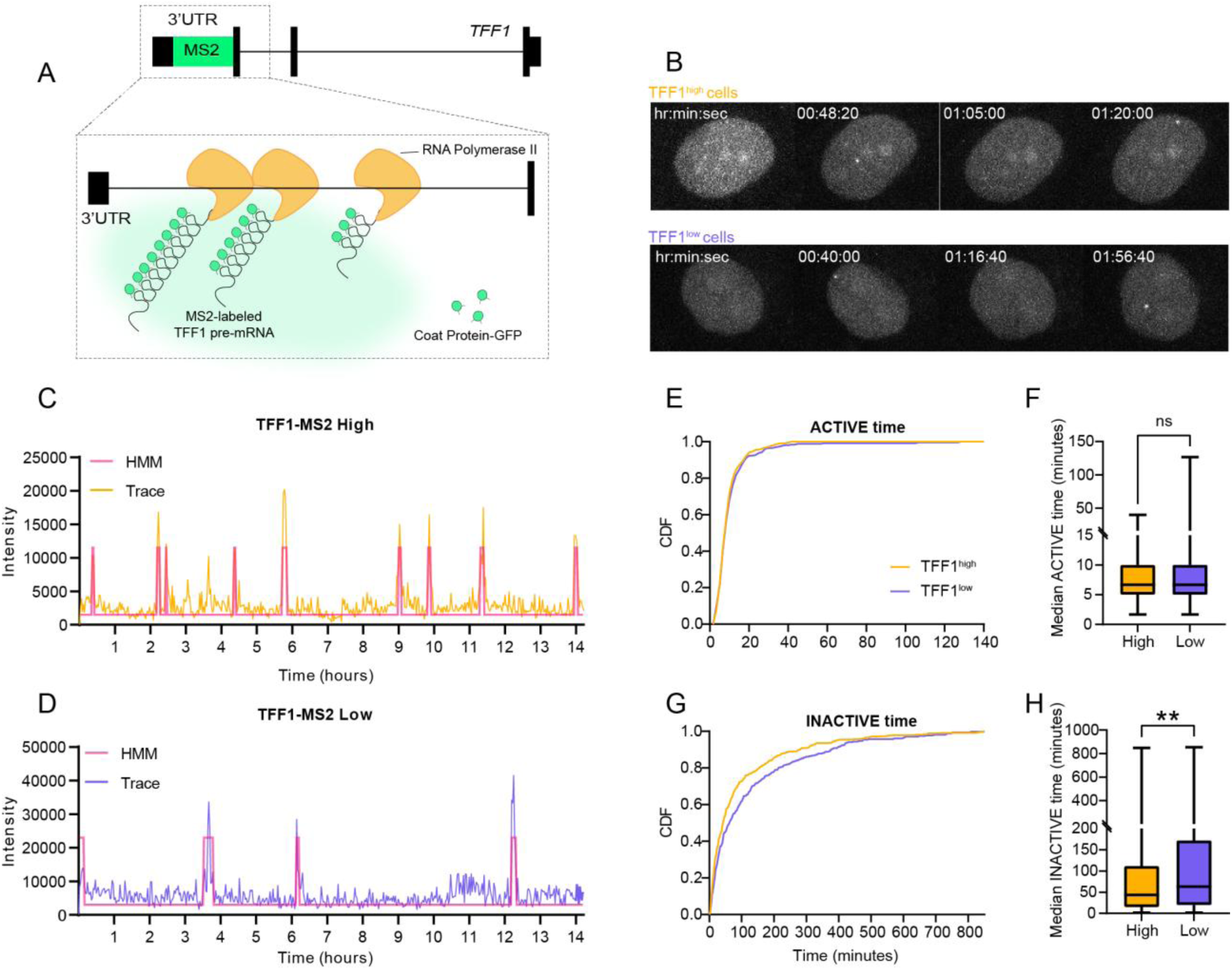
Live-cell imaging of TFF1 transcription reveals that TFF1^high^ cells are hyper-bursting. TFF1-MS2 MCF7 cells were sorted as described before. Sorted high and low TFF1-MS2 cells were maintained overnight after sorting with 1 nM E2 and imaged for 14.2 hours. A. Schematic of the TFF1-MS2 system for live-cell imaging of TFF1 transcription. B. Movie stills of transcribing TFF1^high^ and TFF1^low^ cells. C. Traces of transcription sites tracked in high TFF1 protein expressing TFF1-MS2 cells with HMM calling of active periods. D. Traces of transcription site tracked in low TFF1 protein expressing TFF1-MS2 cells with HMM calling of active periods. E. Cumulative distribution of active periods for both high and low TFF1-MS2 cell traces. F. Box plot of all active times for TFF1^high^ and TFF1^low^ TFF1-MS2 cells. G. Cumulative distribution of inactive periods for both high and low TFF1-MS2 cell traces. H. Box plot of inactive times for TFF1^high^ and TFF1^low^ TFF1-MS2 cells.

### TFF1^high^ and TFF1^low^ cells maintain stable TFF1 transcriptional states

Determining the stability and dynamics of the active and repressive states is critical to developing more effective therapeutic strategies for heterogeneous diseases such as cancer. Given that H3K27me3 is a stable repressive mark associated with prolonged transcriptional repression^29,30^, and its presence across the TFF1 locus in TFF1^low^ cells, we sought to determine the temporal stability of TFF1 expression states. Therefore, we profiled TFF1 expression in sorted cells by single molecule RNA FISH (smFISH) at different timepoints after seeding cells. Sorted TFF1^high^ and TFF1^low^ cells were seeded on glass cover slips and maintained in culture with 1 nM E2 for 4, 6, and 10 days. We visualized TFF1 transcription sites by hybridizing for both the TFF1 exon and intron^12^ (Figure 4a). Individual nuclear intron spots colocalizing with an exon spot were identified as transcription sites (TS) (Figure 4a). We determined the percentage of TFF1 transcribing cells, containing 1 or more active TS, within both TFF1^high^ and TFF1^low^ populations. We observed that TFF1^high^ cells remained significantly more transcriptionally active than TFF1^low^ cells over the time course. Notably, the percentage of TFF1 transcribing cells significantly increased to 30% on Day 6 post-sorting in TFF1^high^ cells, compared to 13% in TFF1^low^ cells (Figure 4b). By day 10, however, both populations showed a tendency to converge toward similar proportions of transcribing cells. This trend was also confirmed at the mRNA level by counting the TFF1 mRNA counts per cell in both populations (Supp. Figure 3a).

**Figure 4.**
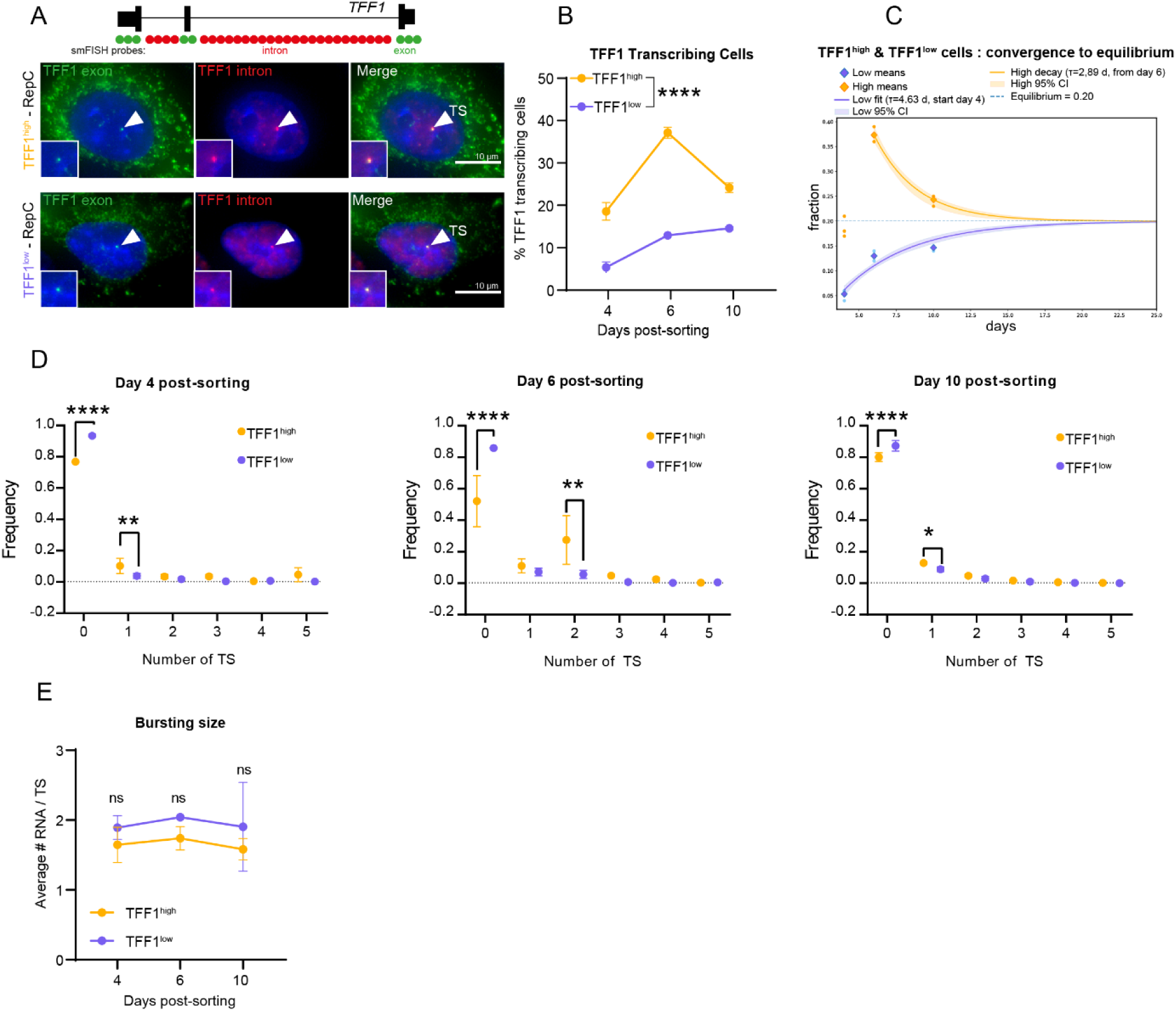
TFF1^high^ and TFF1^low^ cells maintain stable TFF1 transcriptional states. A. Illustration of TFF1 exon (green) and intron (red) smFISH probe locations (Top). Images of TFF1 exon and TFF1 intron smFISH signals (bottom). Colocalized nuclear spots indicate a transcription site (TS, arrow) in TFF1^high^ and TFF1^low^ cells. B. Quantification of the percent of TFF1 transcribing cells in TFF1^high^ and TFF1^low^ cells at days 4, 6 and 10 post sorting. Cells with one or more TS are classified as transcribing cells. C. Equilibrium model reveals that the percentage of TFF1 transcribing cells from TFF1^high^ and TFF1^low^ cell populations converges to 20% after 20 days. D. Individual frequency dotplots of the number of TS per cell in TFF1^high^ and TFF1^low^ cells at days 4, 6 and 10 post sorting. E. Quantification of the number of RNA per TS (burst size) is plotted across the time course in TFF1^high^ and TFF1^low^ cells.

To estimate the time required for re-establishment of the transcriptional equilibrium (≈20% transcribing cells in the bulk population; Supp. Figure 3b), we jointly fit a two-state model to the TFF1^high^and TFF1^low^ transcriptional trajectories. The model assumes that cells are either transcriptionally active or inactive. TFF1^high^ cells represent a population subset enriched for active alleles, whereas TFF1^low^ cells represent a population subset enriched for repressed alleles. Thus, by tracking the return of sorted populations to equilibrium over these timescales, we effectively measure the transition rates into and out of the deep repressive state. Using this approach, the model predicted convergence to equilibrium at ∼21 days post-sorting (Figure 4c). Notably, cells with TFF1 in the deep repressive state required ∼14 days to increase from 6% transcribing cells to the 20% equilibrium, consistent with previous modeling of the deep repression kinetics in bulk cell populations^12^. In addition, rather than assuming a fixed equilibrium at 20% of transcribing cells, we next used the model to fit this parameter. Strikingly, the model predicted an equilibrium fraction *A_eq_* of 17.9%, confirming the observed measurements^31^ (Supp. Figure 3c). Importantly, the observed rate from the deep repressive state to an active state *K_on_* was 0.052 days^-1^ or 19.2 days, which is consistent with the rates observed from bulk cell data^12^, Table S1. We also observed that although TFF1^high^ and TFF1^low^ cells maintained their respective expression states, both populations displayed dynamic temporal behavior over time (Supp. Figure 3d).

We next determined if there were differences in the number of co-active alleles between sorted cell groups, we quantified the number of active alleles at every time point after sorting using smFISH and evaluated the percentage of the cell population exhibiting 0, 1, 2, 3, 4 or 5 TSs (Figure 4d). Strikingly, we observed that at every time point after sorting, TFF1^high^ cell population systematically exhibited a higher percentage of cells with 1 or more active TS than TFF1^low^ cells. These data indicate that more TFF1 alleles in TFF1^high^ cells are stably active for at least 10 days post-sorting than in TFF1^low^ cells.

Finally, we sought to determine if more polymerases were initiated during transcriptional bursts in TFF1^high^ expressing cells than in TFF1^low^ cells by measuring the number of RNA molecules synthesized in each transcription burst (Figure 4e). We divided the TFF1 exon smFISH signal intensity at individual TS by the median intensity of the cytoplasmic spots that correspond to single TFF1 mRNA molecules. We observed that each TFF1 TS contained approximately 2 RNA molecules. Furthermore, this burst size did not change over time in either TFF1^high^ or TFF1^low^ cells (Figure 4e). Taken together, our data demonstrate that the H3K27me3 enriched deep repressive state at the TFF1 locus is stable over extended timescales, maintaining transcriptional silencing for about 19 days. In contrast, TFF1^high^ cells exhibit dynamic transcriptional activity, sustaining high expression and an increased number of active alleles over the same period. Finally, the convergence of the sorted populations to an equilibrium demonstrates that TFF1 alleles dynamically switch between deep repression and activity.

### TFF1 heterogeneity reflects phenotypic heterogeneity associated with endocrine resistance

In ERα+ BC patients, high TFF1 expression is associated with better prognosis^13,14^. This led us to ask whether, in our cell system, TFF1^low^ cells were enriched in a cell state that would constitute an advantage for tumor cells in the context of disease progression. To determine if these cell populations was enriched in specific gene pathways, we assessed the transcriptomic profile of TFF1^high^ and TFF1^low^ populations by combining 2 independent sets of Nascent-seq experiments and by performing differential gene expression analysis using DESeq2 analysis (Supp. Figure 4a,b). 1089 DEGs were identified, including TFF1. We next performed Gene Ontology (GO) analysis using Enrichr^32–34^ for “Molecular Signature Hallmarks”. GO profiling revealed that early and late estrogen response ontology categories were enriched in TFF1^high^ cells. This enrichment consists of a small group of 47 estrogen-responsive genes in high TFF1 cells (Log_2_FC = 1.04) (Figure 5a; Supp. Figure 4c). Enrichr analysis of the TFF1^low^ cells revealed enrichment for genes involved in oncogenic pathways such as STAT signaling, EMT, UV response, Hypoxia, Hedgehog signaling, and angiogenesis (Figure 5b).

**Figure 5.**
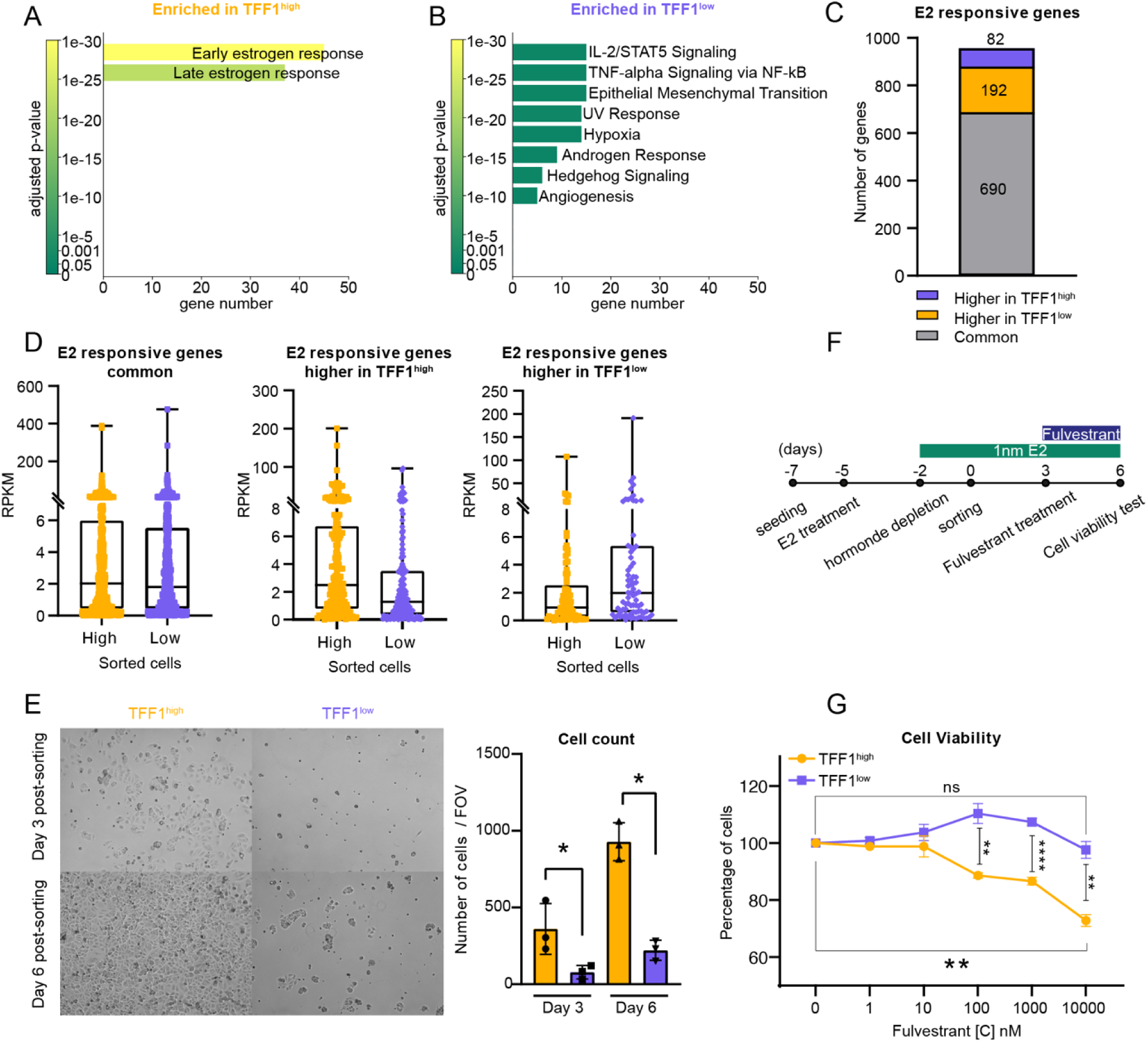
TFF1 heterogeneity reflects phenotypic heterogeneity associated with endocrine resistance. A. EnrichR gene ontology results using Hallmark Msig_Database for genes enriched in TFF1^high^ cells. B. EnrichR gene ontology results using Hallmark Msig_Database for genes enriched in TFF1^low^ cells. C. Overlap of E2 responsive genes with TFF1^high^ and TFF1^low^ RNAseq DEGs. D. Boxplot of E2 responsive genes expression common for TFF1^high^ and TFF1^low^ cells (left), higher TFF1^high^ cells (middle) and higher in TFF1^low^ cells (right). E. Left: Brightfield images of TFF1^high^ and TFF1^low^ cells at days 3 and 6 post sorting. Right: Quantitation of the number of cells in brightfield images at days 3 and 6 post sorting. F. Schematic of cell viability assay design in response to fulvestrant treatment. G. Cell viability quantitation for TFF1^high^ and TFF1^low^ cells in response to different doses of fulvestrant.

We next determined if the enrichment of estrogen-responsive genes in TFF1^high^ and TFF1^low^ cells was due to a significant difference in estrogen responsiveness. To assess the estrogen responsiveness of TFF1^high^ and TFF1^low^ cell populations, we performed RNAseq from TFF1^high^ and TFF1^low^ sorted E2 treated cells and differential gene expression analysis using DESeq2 (Supp. Figure 4d). We identified 1913 DEGs between TFF1^high^ and TFF1^low^ cells that we compared to a list of 964 genes upregulated in MCF7 cells in response to E2 treatment^31^. Depending to their overlap between the two data sets, E2 responsive genes were then classified as significantly “Higher in TFF1^high^”, “Higher in TFF1^low^”, or as “Common” in both TFF1^high^ and TFF1^low^ populations if there was no significant difference. We identified 82 E2-responsive genes upregulated in TFF1^low^ cells, 192 E2-responsive genes upregulated in TFF1^high^ cells, and 690 E2-responsive genes expressed in both cell populations (Figure 4c, Supp. Figure 4e). We plotted the mean RPKM value of each individual E2-responsive gene in TFF1^high^ and TFF1^low^ cells, for each of the identified subsets of genes (Figure 4d). The median expression for common E2 responsive genes was 2 RPKM and 1.99 RPKM in TFF1^high^ and TFF1^low^ samples, respectively. Given that 98% of the 22,929 ER peaks are the same between TFF1^high^ and TFF1^low^ cells (Figure 2), these data suggest that both cell populations retain E2 responsiveness.

Given that TFF1^low^ cells are E2 responsive and are enriched in genes associated with endocrine therapy resistance, such as JAG1^35^, EGFR^10^, VEGF^36^, we tested if this expression heterogeneity could provide benefits to the cell population. We first asked whether TFF1^low^ cells could show proliferative advantages compared to TFF1^high^ cells. After sorting, we plated TFF1^high^ and TFF1^low^ cells at the same cell density and monitored their growth over time (Figure 5e). Unexpectedly, at Day 3 and Day 6 post-sorting, TFF1^low^ cells showed a significantly lower proliferation than did the TFF1^high^ population. We next tested their viability when sorted cell populations were treated with Fulvestrant. Fulvestrant is a selective estrogen receptor degrader (SERD) widely used as standard care endocrine therapy for hormone receptor-positive breast cancers^37,38^. After sorting, the cells were cultured for 3 days in 1nM E2 and subjected to a dose response of fulvestrant for 3 additional days. At Day 6 post-sorting, the cell viability was assessed in TFF1^high^ and TFF1^low^ cells (Figure 5f) using the CellTiter-Glo assay. Strikingly, we observed that the TFF1^low^ cells were more resistant to the increasing dose of fulvestrant than were TFF1^high^ cells, with TFF1^high^ cells showing a dose-dependent decrease in cell viability (Figure 5g). In summary, our results suggest that TFF1^high^ and TFF1^low^ cells remain estrogen-responsive yet exhibit different subsets of estrogen-responsive genes. Moreover, TFF1 heterogeneity is directly associated with a differential response to endocrine therapy.

### Promoters with heterogeneous H3K27me3 deposition are enriched in estrogen-responsive genes and reflect variable gene expression

Our analysis revealed that TFF1 exhibits variable H3K27me3 chromatin and nascent RNA expression across distinct cell populations. Consistent with this, TFF1 expression is heterogeneous in unsorted MCF7 cells^12^. To identify other genes with similar characteristics to TFF1, we first assigned gene symbols to the 2168 TSSs exhibiting differential H3K27me3 deposition identified from our CUT&Tag data (Figure1) and annotated 934 unique genes (representing 8.63% of all expressed genes) (Figure 6a). Gene ontology analysis of those genes using Enrichr revealed several significantly enriched categories, notably including categories of genes related to H3K27me3 signaling in MCF7 cells (Supp. Figure 5a), and in PRC2 components identified as chromatin-bound in MESC cells, such as SUZ12, EZH2, JARID2, and MTF2 (Supp. Figure 5b). Interestingly, we also observed enrichment for the Estrogen Response Early and Late gene categories (Supp. Figure 5c).

**Figure 6.**
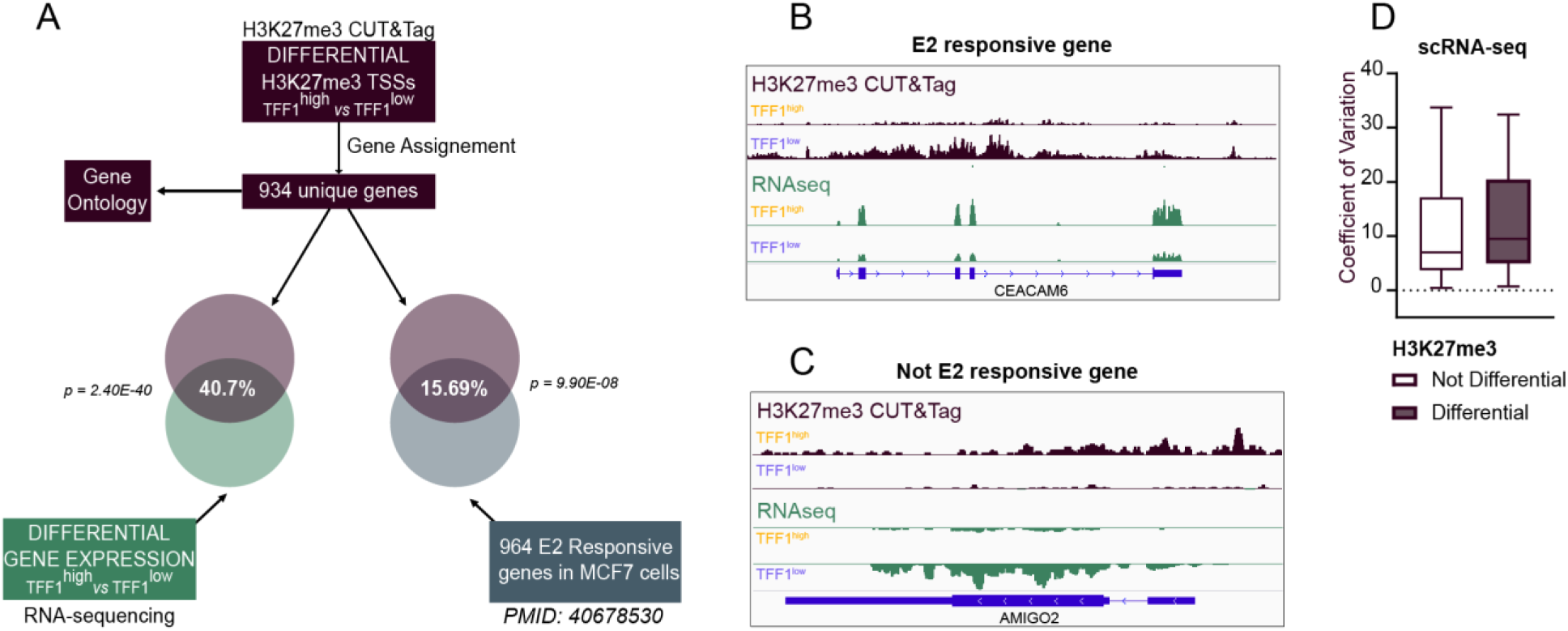
Promoters with heterogenous H3K27me3 deposition are enriched in estrogen responsive genes and are more variably expressed. A. Schematic of analysis of differential H3K27me3 deposition and differentially expressed genes in TFF1^high^ and TFF1^low^ cells. Same analysis was performed for estrogen responsive genes. B. Genome browser view of Ceacam6, an example of estrogen responsive gene with differential H3K27me3 and RNA expression in TFF1^high^ and TFF1^low^ cells. C. Genome browser view of Amigo2, an example of non-E2 responsive gene with differential H3K27me3 and RNA expression in TFF1^high^ and TFF1^low^ cells. D. Boxplot of the Cofficient of Variation for promoters with differential and not differential H3K27me3 deposition.

We next asked if the differential H3K27me3 deposition at these genes predicted a differential expression response. To do so, we integrated the H3K27me3 CUT&Tag data with RNAseq data we obtained from TFF1^high^ and TFF1^low^ sorted populations. From the RNAseq, we observed 1913 DEGs out of 19,703 expressed genes between TFF1^high^ and TFF1^low^ cells. When overlapping genes from the differential analyses of both CUT&Tag and RNAseq datasets, we observed a striking 40.7% overlap, corresponding to 380 genes (Figure 6a, p= 2.4E-40), indicating that differential H3K27me3 deposition at the TSSs correlates with differential expression of those genes.

Given that we observed gene ontology enrichment of the estrogen response in the differential H3K27me3 genes (Supp. Figure 5c), we next asked whether genes with differential H3K27me3 deposition were also enriched in E2 responsive genes. To do so, we compared those genes with the list of E2 responsive genes described above^31^ in Figure 5. Strikingly, we observed that 15.69% of them were identified as E2 responsive, which represent a significant enrichment (p = 9.90e-08) (Figure 6a). We illustrate two E2-responsive genes, CEACAM6 and BAMBI (Figure 6b; Supp. Figure 6d) and two non E2-responsive genes, AMIGO2 and c14orf132 (Figure 6c; Supp. Figure 6e), with differential H3K27me3 deposition (CUT&Tag, purple), and differential RNA expression (RNAseq, green).

Lastly, we asked if the 934 genes with differential H3K27me3 deposition were more variably expressed than the genes without variable H3K27me3. Using publicly available MCF7 scRNA-seq data^39^, we calculated the coefficient of variation across cells. This dataset is a timeseries after induction with 100 nM of E2. We first identified DEGs at the time point 24H and calculated the coefficient of variation for all genes, and observed that genes with differential H3K27me3 were more variably expressed than were genes without differential H3K27me3 (Figure 6d, p=0.0028, t-test). Taken together, we discovered that TFF1 expression heterogeneity in the cell population reflected a broader heterogeneity in chromatin landscape, some of which is marked by H3K27me3. Moreover, our data suggests that differential H3K27me3 is a general mechanism that regulates variable gene expression and heterogeneous phenotypes.

## Discussion

Gene expression heterogeneity within cancer cell populations presents a formidable barrier to effective treatment, enabling dynamic adaptation and resistance to therapy^40^. This heterogeneity is well documented in ERα⁺ breast cancer, yet its epigenetic basis remained uncharacterized^41^. Our study revealed that focal H3K27me3 deposition, a hallmark of Polycomb Repressive Complex 2 (PRC2) activity^17^, plays a central role in establishing heterogeneous expression of the estrogen-responsive gene TFF1. While some cells maintain transcriptionally active, hyper-bursting TFF1 loci, others exhibit prolonged-stable repression accompanied by H3K27me3 enrichment. These chromatin-defined subpopulations are also functionally distinct. TFF1^low^ cells display increased resistance to the estrogen receptor antagonist fulvestrant in comparison to TFF1^high^ cells. Importantly, this transcriptional heterogeneity does not arise from differential ERα availability or global binding differences. It is a result of localized epigenetic restriction of receptor access. Extending beyond TFF1, our genome-wide H3K27me3 profiling revealed an enrichment of differentially marked estrogen-responsive genes that show increased cell-to-cell expression variability. Our work supports a broader role for Polycomb in regulating transcriptional heterogeneity across the estrogen signaling network. Whether H3K27me3 deposition in mammals is deterministic or subject to stochastic, environmentally-sensitive^42^ regulation remains an open question. Our data highlight the critical role of chromatin context in shaping estrogen responsiveness and points to a limitation of therapies that target ERα alone.

Polycomb group (PcG) proteins, primarily acting through the Polycomb Repressive Complex 2 (PRC2), have long been viewed as key regulators of developmental gene silencing^17,43^. However, our data support an expanded role for Polycomb in breast cancer. H3K27me3 chromatin regulates heterogeneous and possibly reversible repression. Rather than uniformly silencing genes across the cell population, Polycomb functions as a stochastic gatekeeper of transcriptional accessibility^44^. It regulates how many alleles per cell or cells in a population remain in a transcriptionally permissive state. In this model, single aneuploid MCF7 cells may have between 0 and 5 alleles marked by H3K27me3. Interestingly, the other PcG complex, PRC1, can also regulate the number of alleles that are switched ON^45^. This probabilistic mode of repression suggests a plastic epigenetic landscape in which Polycomb-mediated silencing contributes to both phenotypic diversity and therapeutic resistance. Notably, the deeply repressed TFF1 state persists for days, even under estrogen stimulation. This is consistent with a form of epigenetic memory^46^. Moreover, prior work shows that TFF1 alleles can transition into and out of this repressive state^12^. This flexibility may allow tumors to generate subpopulations that are temporarily insulated from treatment pressure.

Indeed, our findings reveal that TFF1^low^ cells are more resistant to fulvestrant. However, they proliferate significantly less than TFF1^high^ cells under estrogenic conditions. TFF1^low^ cells exhibit similar expression of the majority of ER targets as TFF1^high^ cells, and reduced expression of a smaller group of ER targets. These results suggest that estrogen signaling persists in these cells but is uncoupled from proliferative programs. This phenotype may reflect a form of Polycomb-regulated dormancy, in which cells retain estrogen signaling but are epigenetically insulated from apoptosis. This state resembles drug-tolerant persister cells seen in other cancers, which evade treatment through quiescence rather than intrinsic resistance^47^. Therefore, TFF1^low^ cells may not be intrinsically resistant, but rather evade cytotoxicity through slow cycling and reduced ERα turnover. These are all hallmarks of residual disease in cancer^47^.

In summary, our data suggest that Polycomb and H3K27me3-marked chromatin may facilitate an epigenetic bet-hedging strategy, enabling survival and adaptation through dynamic tuning of gene promoter accessibility. Interestingly, correlations between heterogeneous H3K27me3 chromatin and cancer phenotypes have been observed in head and neck carcinomas^48^, and in leukemias^49^. Together these studies indicate a more general mechanism of therapeutic resistance in cancers regulated by Polycomb. These insights carry important therapeutic implications. Targeting PRC2 with EZH2 inhibitors could restore expression of silenced genes and sensitize resistant subpopulations. Indeed, targeting of the histone demethylase KDM5B with small molecule inhibitors led to reduced heterogeneity and increased sensitivity to an estrogen receptor antagonist^50^. However, such interventions are complicated by the roles of Polycomb complex members in both repression and activation^21,51^. Future strategies may require more precise tools capable of disrupting repressive Polycomb functions without impairing its regulatory roles in gene activation. Ultimately, understanding how Polycomb interacts with transcriptional bursting, enhancer dynamics, and hormonal signaling over time may reveal new avenues for reducing heterogeneity and overcoming resistance in hormone-responsive cancers.

### Authors’ contributions

Conceptualization and formal analysis: FC, JR. Investigation: FC, CD, LK, PY, CB. Methodology: FC, JR. Bioinformatics: BB, CD, JR. Microscopy: FC, ES, CT. Visualization: FC. Writing: FC, JR.

## Acknowledgments

We would like to thank Dr. Negin Martin at the Viral Vector Core Facility for generating the lentiviruses; Dr. Guang Hu, Dr. Xin Xu, Jason Malphurs, Jason Li and Gregory Solomon for providing Next Generation Sequencing services. We would like to thank Donna Stefanick and Dr. Carl Bortner for Flow Cytometry services. We also would like to thank Dr. Paul Wade, Dr. Trevor Archer, Dr. Dave Levens and Dr. Jackson Hoffman for fruitful discussion regarding this study. This research was supported by the Intramural Research Program of the NIH (ES103331).

## Methods

### Cell Culture

MCF7 cells (ATCC) were cultured in MEM supplemented with 10% FBS (Sigma), 1% Glutamine and 1% Penicillin/streptavidin (Gibco). For estrogen induction experiments, MCF7 cells were plated in MEM and allowed to recover for 48 hours before being hormone depleted. For hormone depletion (HD), cells were washed 2 times with 1X PBS and incubated 1 hour at 37°C with phenol red-free MEM supplemented with 5% charcoal-stripped FBS (CS-FBS), (HD medium). This step was repeated one more time. After 72 hours of HD, cells were stimulated with 1 nM of 17β-estradiol (E2) or 1 nM vehicle (ethanol, OH) during 48 hours.

### Single molecule RNA FISH and ImmunoFISH

MCF7 cells were plated on 18 mm No. 1.5 coverslips in 12 well-plates and HD as previously described. After 48 hours of E2 stimulation, cells were washed 2 times with 1X PBS, fixed with 4% PFA for 10 min at room temperature, then washed 2 times with PBS and incubated overnight at 4°C in 70% ethanol for membrane permeabilization. The day after, smFISH was performed following the adherent mammalian cell Stellaris RNA FISH Protocol. For immunoFISH, an immunostaining was performed prior to the smFISH protocol. Briefly, cells were incubated with the primary antibody (Table S2) overnight at 4°C, then wash 3 times with 1X PBS and incubated 1 hour with the secondary antibody at room temperature, in the dark. After the secondary antibody staining, cells were washed 3 times with 1X PBS and fixed with 4% PFA for 10 min at room temperature. After 2 washes with 1X PBS, smFISH protocol was performed. Custom Stellaris™ RNA FISH Probes were designed utilizing the Stellaris RNA FISH Probe Designer using a masking level of 5, oligo length of 20, minimum spacing of 2 nucleotides (LGC, Biosearch Technologies, Petaluma, CA) available online at www.biosearchtech.com/stellarisdesigner. The cells were hybridized with the Stellaris RNA FISH Probe set labelled with Quasar™ 570 dye or Quasar™ 670 dye (Biosearch Technologies), following the manufacturer’s instructions available online at www.biosearchtech.com/stellarisprotocols (Table S2).

### Microscopy

(Immuno)FISH images were acquired using a custom widefield microscope. This microscope uses an ASI (www.asiimaging.com/) Rapid Automated Modular Microscope System (RAMM) base, an ASI MS-2000 Small XY stage, a Hamamatsu ORCA-Flash4 V3 CMOS camera (https://www.hamamatsu.com/, C13440-20CU), an ASI High Speed Filter Wheel (FW-1000), ASI MS-2000 Small XY stage, and a Zeiss C-Apochromat 40x /1.20 NA UVVIS-IR objective. DAPI, Quasar 570 (Cy3), Quasar 670 (Cy5), and 750 were excited using a Lumencore SpectraX (https://lumencor.com/) with violet, green, red and far-red filters. The microscope was controlled using Micro-Manager^52^. 25 fields of view consisting of a 10-micron Z stack with 0.5-micron intervals were acquired per sample. Maximum intensity projections were processed using Fiji and used as input for analysis.

Live-cell imaging was performed using Andor Dragonfly Spinning Disk Confocal (https://andor.oxinst.com/) at 37°C and 5% CO2. 12 to 18 fields of view were imaged using 488 nm excitation, on a 60 X objective with a pinhole size of 40 μm, for 200 msec exposure. Total nuclear volume was imaged using 11 Z planes with 0.8-micron interval. Images were acquired every 100 sec for 512 repeats (14.2 hours). Analysis was performed using maximum intensity projections performed on FIJI (https://fiji.sc).

### Image analysis

(Immuno)FISH data (maximum intensity projections) was analyzed by using custom python scripts adapted^31^. smFISH spots were called by fitting a 2D Gaussian mask and performing local background subtraction. Masks of the cytoplasm and nuclei were generated using Cellprofiler (https://cellprofiler.org/). Transcriptions sites (TS) were identified as overlapping nuclear intron and exon smFISH spots. Overlap was calculated using the Euclidean distance of less than or equal to 5 pixels. The number of nascent transcripts at the TS was determined on a per cell basis by extracting the intensity of the exon signal at the identified TS and dividing by median intensity. Only TS with a minimum of at least 5 exon smFISH spots detected in the cytoplasmic mask were considered in the burst size calculation.

### Cell Sorting

Parental or TFF1-MS2 MCF7 cells were seeded in MEM and allowed to recover for 48 hours before HD. After 72 hours in HD medium, cells were treated with 1 nM E2 for 48 hours. Cells were then incubated 3 min at 37°C with 3 mL of TrypLE before reaction was stopped with 3 mL of prewarmed HD medium. Cells were scrapped and passed 15 times through a 19 G needle before being pelleted at 1,500 rpm for 5 min. This step was repeated one more time to obtain single cell suspension and avoid cell clumping. 10 to 15 million cells were further blocked 15 min in blocking buffer (10% CS-FBS, 250 μL EDTA, 1% BSA, in 1X PBS) and then incubated with anti-TFF1 antibody, 1/100 dilution (#MA5-32781, Thermo) for 1 hour on a nutator at 4°C in staining buffer (3% CF-FBS, 250 μL EDTA, 1% BSA, in 1X PBS), followed by 1 hour incubation on a nutator at 4°C in the dark with an anti-Rabbit secondary antibody, Alexa Fluor 647 (#A32733, Invitrogen). Cells were examined using a BD FACSAriaII or a BD Symphony S6 cell sorter (Becton Dickinson Biosciences, San Jose, CA) equipped with FACSDiVa software. Initially, a “scatter” gate was set on a forward scatter (FSC-A) versus side scatter (SSC-A) dot plot to isolate the principal population of cells free of debris. Subsequently, cells were consecutively gated on a side scatter height (SSC-H) versus width (SSC-W), then a forward scatter height (FSC-H) versus width (FSC-W) dot plot to isolate single cells. Dead cells were excluded using PI (Ex: 561; Em: 585) on a SSC -A versus PI dot plot. Cells stained with APC (Ex: 633; Em: 660) were examined on a FSC-A versus APC-A dot plot. The top 20% of APC expressing cells were sorted as “TFF1^high^” population, and the lower 55% to 70% were also sorted as “TFF1^low^” populations. Cells were then pelleted by centrifugation 20 min at 1,500 rpm, 4°C and further seeded on poly-L-lysine pre-coated coverslips in 5X Pen/Strep supplemented HD medium containing 1 nM E2, or processed for Nascent RNA, total RNA, or nuclear isolation, respectively for Nascent RNAseq, total RNAseq, or CUT&Tag experiments.

### Chromatin-bound RNA extraction (Nascent RNAseq)

Sorted cell pellet were resuspended in hypotonic buffer (10 mM Hepes pH 7.3, 1.5 mM MgCl2, 10 mM KCl. 0.5 mM DTT) and incubated 10 min on ice. Cell lysates were then dounced 2 times using loose then tight pestles and loaded on 20% sucrose cushion in hypotonic buffer. Nuclei were pelleted 10 min at 3,000 rpm at 4°C. The supernatant (cytoplasmic fraction) was discarded and nuclei were washed 3 times using 1X PBS supplemented with 1 mM EDTA and further resuspended in 300 μL of 1X PBS-1 mM EDTA. Chromatin fraction was extracted by drop-by-drop addition of 1 volume of 2X NUN buffer (600 mM NaCl, 2% NP-40, 2 M urea, 2 mM DTT, 40 mM Hepes pH 7.3) while nuclei resuspension was gently mixed on Vortex position 3. After 20 min incubation on ice, chromatin was pelleted 30 min at 14,000 rpm at 4°C. Supernatant (nucleoplasmic fraction) was discarded and the chromatin-associated RNA was further extracted by resuspending the chromatin pellet in 1 mL of Trizol and incubated at 65°C for 15 min or until complete dissolution. Next, chloroform was added and RNA was isopropanol precipitated overnight at -20°C with 1 μL RNA-grade glycogen. The day after, RNA was pelleted for 30 min at 12,000 rpm at 4°C, washed 2 times with 70% Ethanol. RNA pellet was then allowed to dry at room temperature for 10 minutes and resuspended in H2O.

### Total RNA extraction

Sorted cell pellets were resuspended in 1 mL of Trizol and 200 μL of Chloroform. After 15 min centrifugation at 14,000 rpm, the upper phase containing RNA was isopropanol precipitated overnight at -20°C with 1 μL RNA-grade glycogen (Thermo). The day after, RNA was pelleted for 30 min at 12,000 rpm at 4°C, washed 2 times with 70% Ethanol. RNA pellet was then allowed to dry at room temperature for 10 minutes and resuspended in H2O.

### RNA-seq library preparation and analysis

50 ng of total RNA or 2 ng of Nascent RNA were processed using NEBNext Ultra II directional RNA library prep kit for Illumina (#E7760) using section 1 “Protocol for use with NEBNext Poly(A) mRNA Magnetic Isolation Module” or section 4 “Protocol for use with purified mRNA”, respectively. Library profiles were analyzed using TapeStation and material quantified using Qubit. RNA was then sequenced with 75 bp single-end reads on Illumina NextSeq high-output. Reads were filtered so that only those with a mean quality score of 20 or greater were kept. Adapter was trimmed using Cutadapt version 3.7. Reads were aligned to the hg38 genome assembly using STAR version 2.6.0c. Counts were obtained using the featureCounts tool from the Subread package version 1.5.1 with the GENCODE basic gene annotation version 44. For nascent RNA, featureCounts was run using additional parameter “-t transcript” to include reads from introns. Differential expression was quantified using the DESeq2 R package version 1.34.0.

### CUT&Tag library preparation and analysis

CUT&Tag was performed following the Bench top CUT&Tag V.3 protocol^53^. Briefly, 25,000 nuclei were used. 10 µL BioMag-Plus Concanavalin (Bangs Laboratories) were used per reaction to immobilize the nuclei. Primary antibodies were added at dilutions of 1:50 for H3K27ac (#MA5-23516, Invitrogen), for H3K27me3 (#9733, Cell Signaling Technology) and ERα (#8005, Santa Cruz) and incubated for 2 hours. After one hour of secondary antibody incubation, 1.25μL of CUTANA pAG-Tn5 adapter complex was added to load the enzyme to the antibody bound regions. This was followed by one hour of tagmentation at 37°C, after which DNA was extracted using MinElute PCR Purification Kit (Qiagen). Extracted DNA was amplified by PCR using unique primers sets (Nextera XT v2 Full set (N7-S5)) with all reactions undergoing 15 PCR cycles. Libraries were size selected using AMPure XP Beads (Beckman Coulter) with 1.3X ratio. Library quantification and quality assessment were performed using the Qubit Flex and Tapestation. Sequencing was conducted with 50 bp paired-end reads on the Illumina NovaSeq SP high-output platform.

Raw CUT&Tag reads were processed using an adapted pipeline^54^. First, Nextera adaptors were trimmed using Cutadapt before reads were aligned to the hg38 genome assembly using Bowtie2. Reads were sorted using Samtools and deduplication was performed using Picard Tools (http://broadinstitute.github.io/picard). Coverage files were generated using bamCoverage and RPKM normalization. Expressed transcription start sites (TSS) were identified using publicly available start sequencing data^24^. Start-seq data was processed as previously described^24^ but using hg38 and GENCODE v45. Called TSS associated with annotated transcripts were used for calling differential TSS. Differential TSS were identified using a 2kb window surrounding the TSS and reads within the windows were counted using featureCounts. Differential TSS were identified using the DESeq2 R package. ERα heatmaps were made by taking the called peaks and removing peaks overlapping the following exclusion list: https://www.encodeproject.org/files/ENCFF356LFX/. We also excluded a large region on chromosome 21 8205132-8988703 from the upregulated in TFF1high cells which contained very high signal corresponding to very broad peaks. H3K27me3 and ERα peaks were sorted by signal and plotted with deepTools^55^.

### Bioinformatics Analysis of Differential H3K27me3 overlap

*Background gene normalization:* To determine the appropriate background datasets for comparisons, we only analyzed genes considered in both compared datasets. We then performed a hypergeometric test to determine significance. For variable expression analysis, we downloaded the high-quality cell matrix file data from GEO. We calculated the coefficient of variation across cells using the seurat object. We considered only genes that had a calculated coefficient of variation for further analysis. We subsequently binned the gene promoters with differential H3K27me3 and no difference into two groups.

### Relaxation to equilibrium modeling

For modeling of the sorted populations return to equilibrium, cells with transcription sites are denoted as Active and those without transcription sites as Inactive. Cells can transition between these two states :

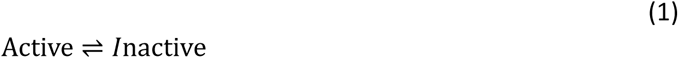

The change in the Active fraction then follows:

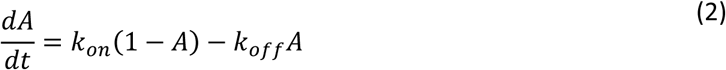

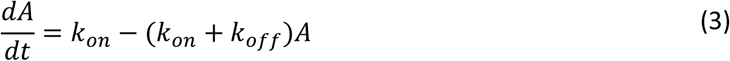

The total relaxation rate to equilibrium 𝜆 :

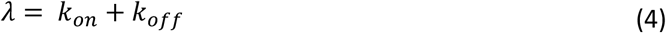

Solving for *A* at equilibrium, equation 3 becomes :

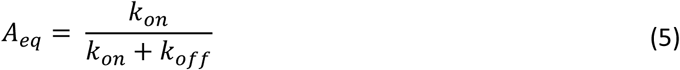

Solving for *k_on_* in terms of *A_eq_* and 𝜆

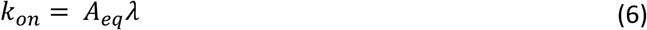

Substituting back in equation 3

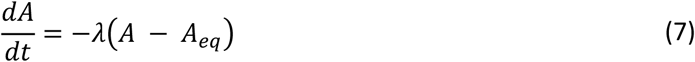

Results in :

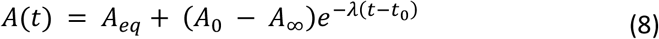

Confidence intervals were calculated by sampling with replacement from the data trajectories.

### GEO Numbers

The data for E2 responsive genes RNAseq was obtained from GEO accession code GSE251654^31^. The scRNAseq was from from GEO accession code GSE154873^39^.

### Enrichr Analysis

Enrichr analysis was done by inputing a gene list into https://maayanlab.cloud/Enrichr/^32–34^

### Statistical Analysis

Statistical tests and p-values are indicated in the figure legends. *P* values ≤ 0.05 were considered significant in GraphPad Prism. All statistical tests were 2-sided.

### Data availability

Sequencing data have been deposited in GEO under accession code GSE280618.

## Supplementary material

**Table S1.** Modeling rates

|  | 1/days | 95CI |
| --- | --- | --- |
| lambda: | 0,290899 | 0.256926 - 0.335398 |
| Aeq: | 0,179115 | 0.171807 - 0.186465 |
| kon: | 0,0521043 | 0.045866 - 0.059653 |
| koff: | 0,238795 | 0.210739 - 0.276273 |

**Supplementary Figure 1.**
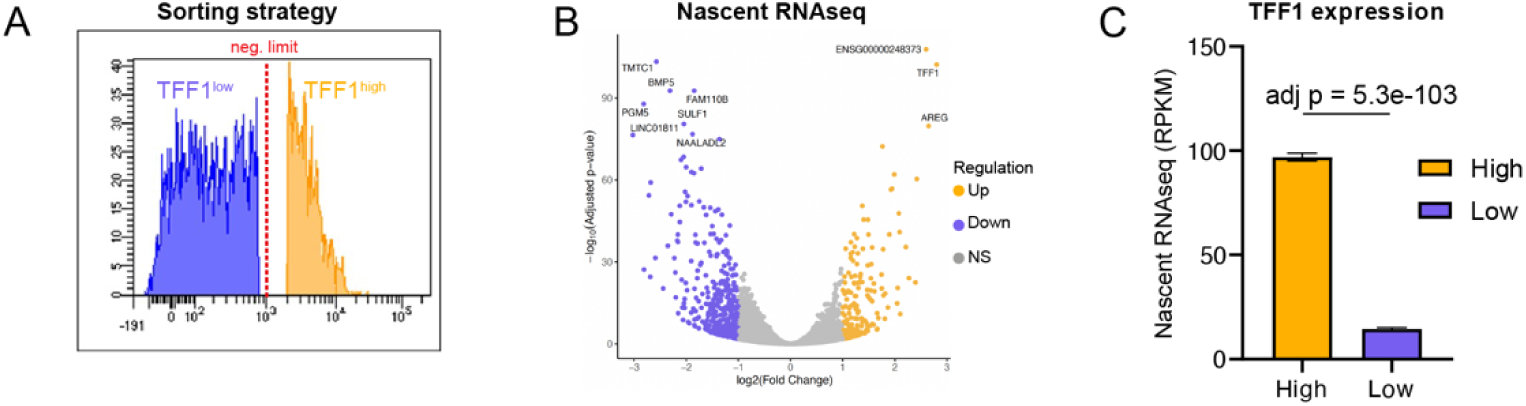
A. Histogram of TFF1 sorting strategy B. Volcano plot of Nascent-RNAseq differentially expressed genes between TFF1^high^ and TFF1^low^ cells. The yellow dots represent significantly upregulated genes, the blue dots represent significantly downregulated genes (log2 FC ≥ 1 and adj p < 0.05), and the grey dots represent insignificant differentially expressed. C. Barplot representing RPKM values of TFF1 expression in Nascent RNAseq in TFF1^high^ compared to TFF1^low^ cells, shows significantly enrichment in TFF1^high^ cells compared to TFF1^low^ cells.

**Supplementary Figure 2.**
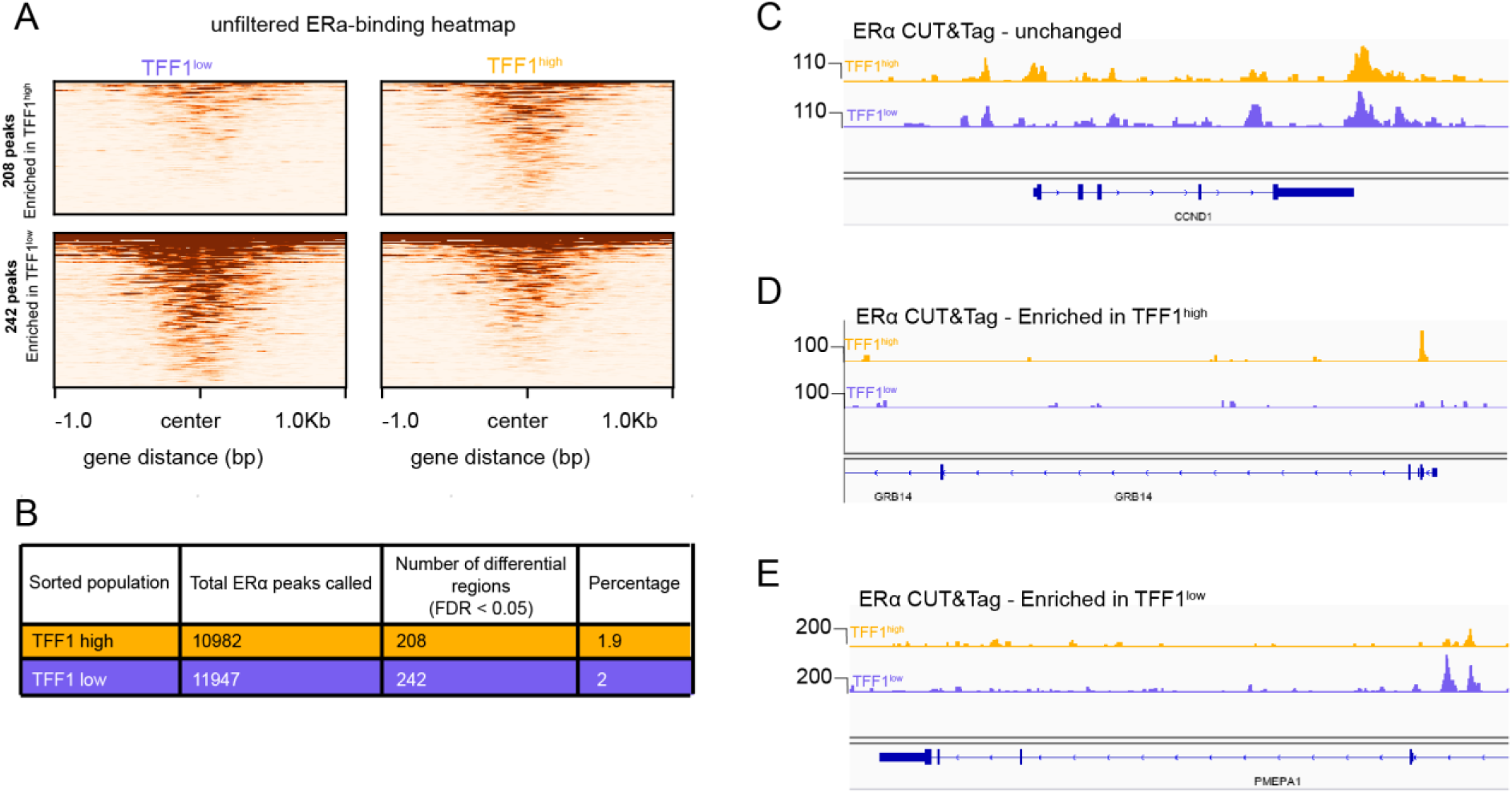
A. Heatmaps of ERa binding in TFF1^high^ and TFF1^low^ cells. B. Table summarizing the total number of ERα CUT&Tag peaks, differentially bound regions and associated percentages of differential regions in TFF1^high^ and TFF1^low^ cells. C. Genome browser view of ERα CUT&Tag shows an example of unchanged ERα binding in TFF1^high^ cells and to TFF1^low^ cells. D. Genome browser view of ERα CUT&Tag shows an example of enriched ERα binding in TFF1^high^ cells compared to TFF1^low^ cells. E. Genome browser view of ERα CUT&Tag shows an example of enriched ERα binding in TFF1^low^ cells compared to TFF1^high^ cells.

**Supplementary Figure 3.**
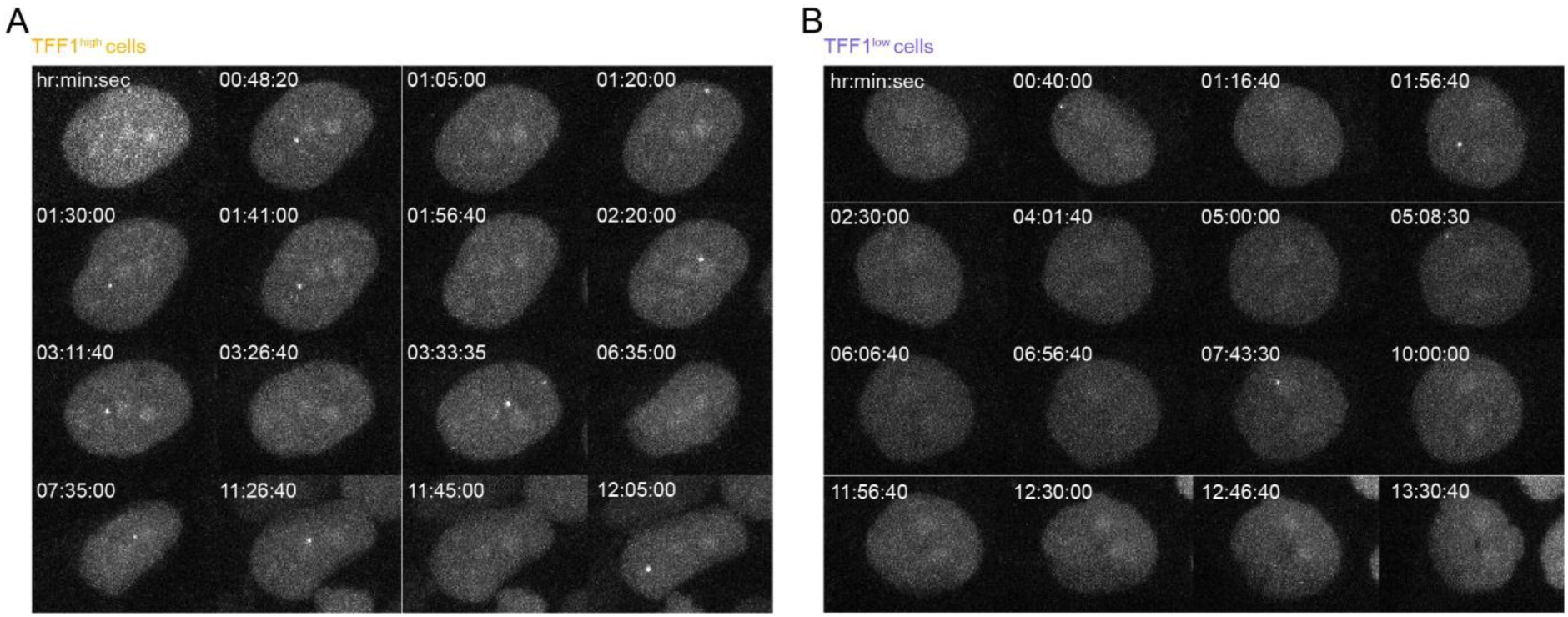
A. Tiles of high TFF1 expressing TFF1-MS2 cells using live-cell imaging after sorting over 14.2 hours. B. Tiles of low TFF1 expressing TFF1-MS2 cells using live-cell imaging after sorting over 14.2 hours.

**Supplementary Figure 4.**
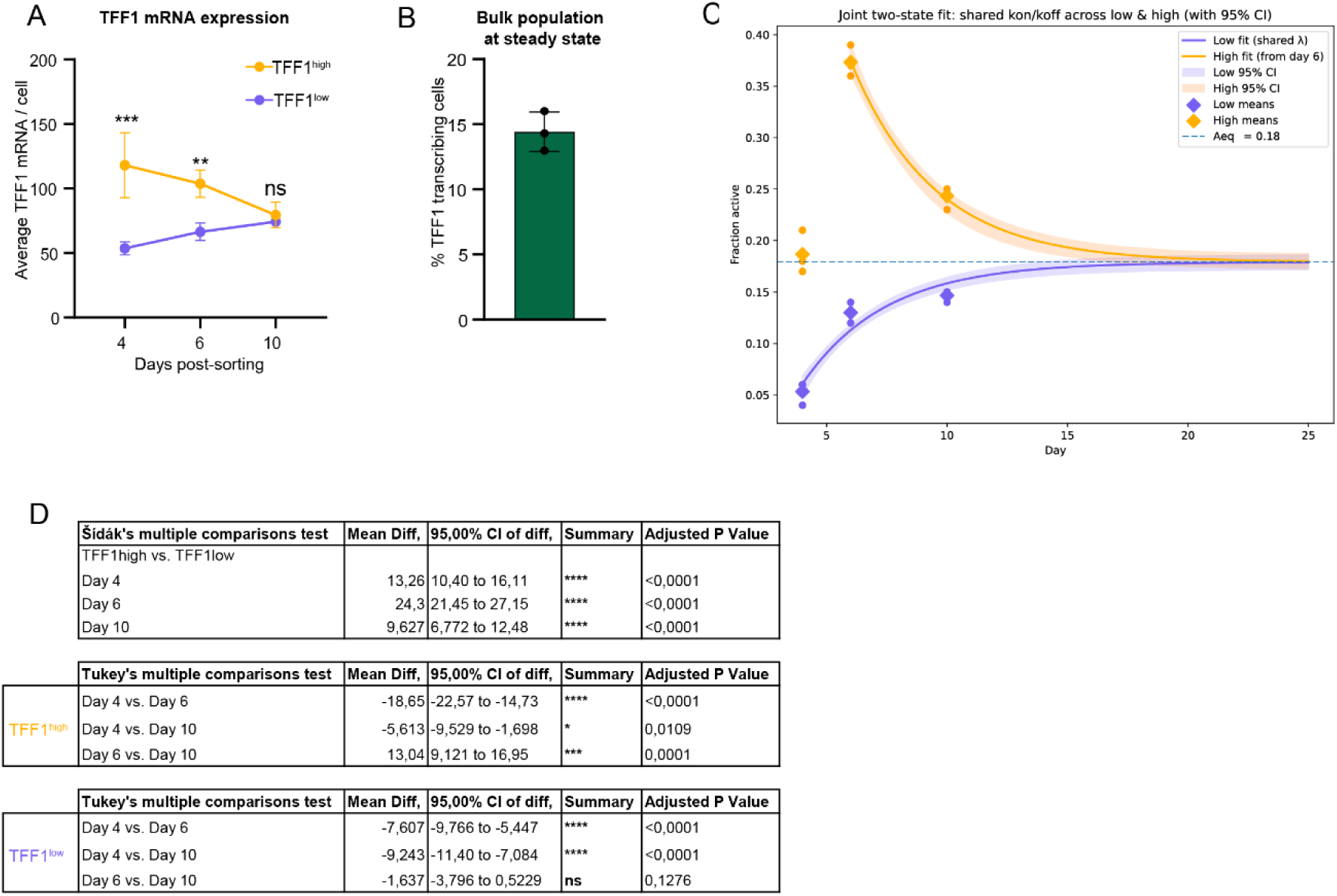
A. Quantification of average number of TFF1 mRNA molecules/cell plotted at days 4, 6 and 10 post sorting in TFF1^high^ and TFF1^low^ cells. B. Percentage of TFF1 transcribing cells in unsorted cell population stimulated with 1nm E2. C. Equilibrium model reveals that the percentage of TFF1 transcribing cells from TFF1^high^ and TFF1^low^ cell populations reach an equilibrium at 17.9% after 20 days. D. Table of statistical analysis showing TFF1 transcription dynamics in TFF1^high^ and TFF1^low^ cells. Comparisons have been made between TFF1^high^ and TFF1^low^ cells over the 10 days-time course after sorting (top table), and between individual days post-sorting (D4 vs D6; D4 vs D10; D6 vs D10) in TFF1^high^ (middle table) and TFF1^low^ cells (bottom table).

**Supplementary Figure 5.**
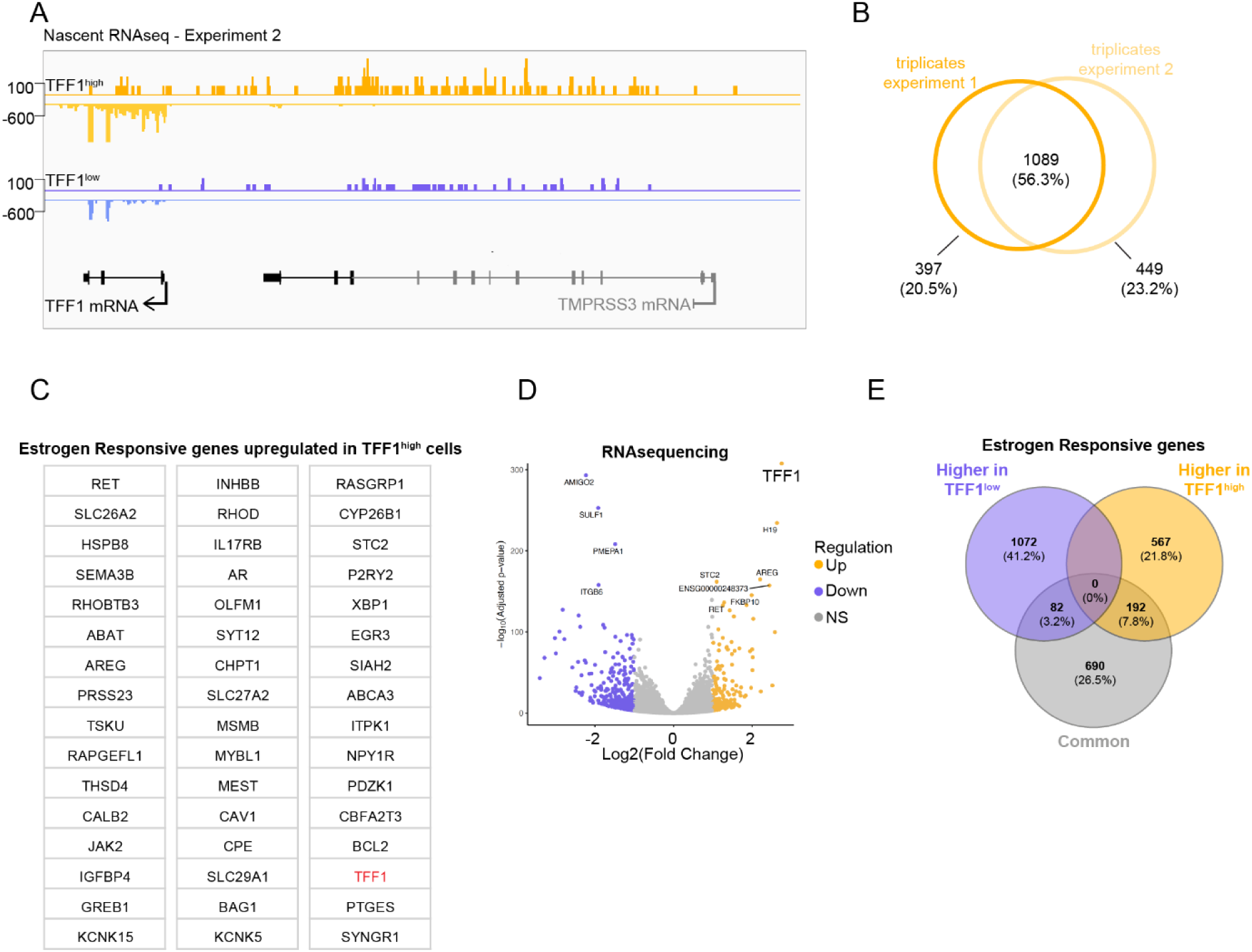
A. Genome browser view of TFF1 locus in another set of Nascent RNAseq (triplicates) confirms that transcription at TFF1 locus in increased in TFF1^high^ compared to TFF1^low^ cells. B. Venn Diagram of overlapping DEGs from the 2 sets of Nascent RNAseq experiments (each set is composed of biological triplicates) shows 1089 common DEGs. Those 1089 DEGs have been used for Gene Ontology analysis. C. Talbe of the subset of 47 estrogen responsive genes upregulated in TFF1^high^ cells. D. Volcano plot of RNAseq DEGs between TFF1^high^ and TFF1^low^ cells. The yellow dots represent significantly upregulated genes, the blue dots represent significantly downregulated genes (log2 FC ≥ 1 and adj p < 0.05), and the grey dots represent insignificant differentially expressed. TFF1 is the most upregulated gene. E. Venn Diagram representing the distribution of estrogen responsive genes enriched in TFF1^high^ cells (yellow), enriched in TFF1^low^ cells (blue), or unchanged between the two cell populations (grey).

**Supplementary Figure 6.**
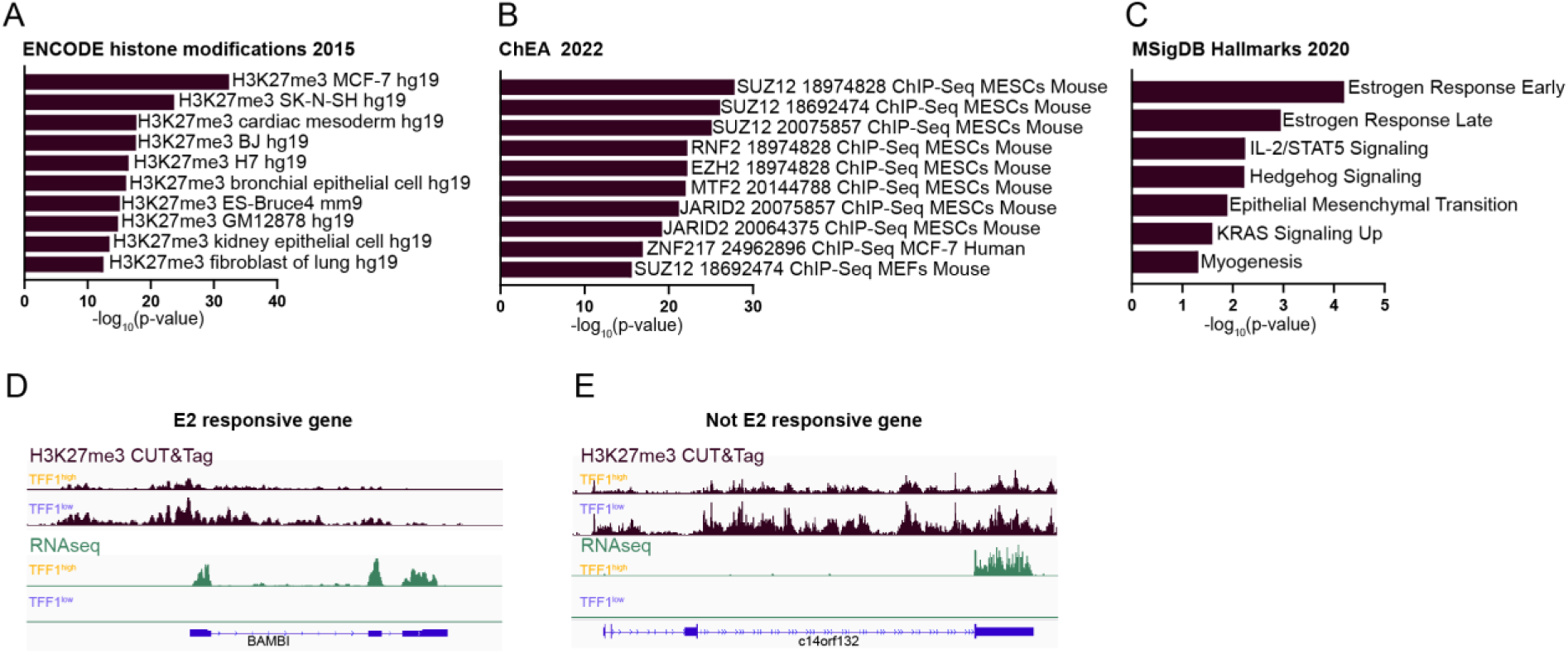
EnrichR gene ontology results for the 934 unique genes with differential H3K27me3 deposition between TFF1^high^ and TFF1^low^ cells using different databases: A. ENCODE Histone Modifications 2015 database shows an enrichment of H3K27me3 deposition on those genes in different cell lines B. ChEA Transcription Factor Targets 2022 database shows an enrichment of Polycomb (PRC2) group members binding to those genes C. Hallmark Msig Database 2020 shows an enrichment of Estrogen Response categories. D. Genome browser view of an example of an E2 responsive gene (BAMBI) showing differential H3K27me3 deposition and differential gene expression in TFF1^high^ and TFF1^low^ cells. E. Genome browser view of an example of a not-E2 responsive gene (c14orf132) showing differential H3K27me3 deposition and differential gene expression in TFF1^high^ and TFF1^low^ cells.

**Table S2.**
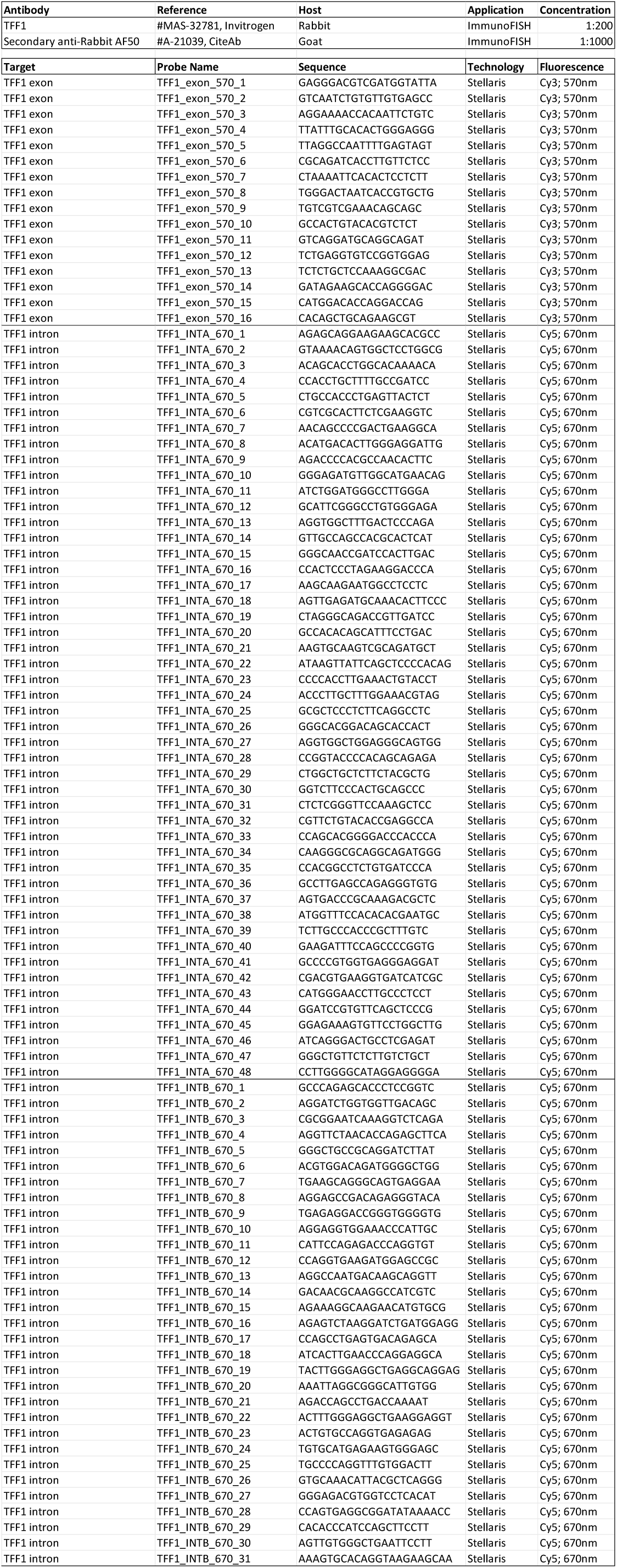
Antibodies and smFISH probes

